# 3D multi-omic mapping of whole nondiseased human fallopian tubes at cellular resolution reveals a large incidence of ovarian cancer precursors

**DOI:** 10.1101/2025.09.21.677628

**Authors:** André Forjaz, Vasco Queiroga, Yajuan Li, Alfredo Hernandez, Ashleigh Crawford, Xiaoyu Qin, Mei Zhong, Margarita Tsapatsis, Saurabh Joshi, Donald Kramer, Oliver Nizet, Habin Bea, Yuhan Li, Shuo Qin, Robert O’Flynn, Mingder Yang, Brittany Pratt, Fan Wu, Paul Gensbigler, Max Blecher, Pei-Hsun Wu, Luciane T. Kagohara, Ie-Ming Shih, David Zwicker, Mark Atkinson, Lingyan Shi, Rong Fan, Ashley L. Kiemen, Denis Wirtz

## Abstract

Uncovering the spatial and molecular landscape of precancerous lesions is essential for developing meaningful cancer prevention and early detection strategies. High-Grade Serous Carcinoma (HGSC), the most lethal gynecologic malignancy, often originates from Serous Tubal Intraepithelial Carcinomas (STICs) in the fallopian tubes, yet their minute size and our historical reliance on standard 2D histology contribute to their underreporting. Here, we present a spatially resolved, multi-omics framework that integrates whole-organ 3D imaging at cellular resolution with targeted proteomic, metabolomic, and transcriptomic profiling to detect and characterize microscopic tubal lesions. Using this platform, we identified a total of 99 STICs and their presumed precursors that harbor TP53 mutations in morphologically unremarkable tubal epithelium in all five specimens obtained from cancer-free organ donors with average-risk of developing ovarian cancer. Although these lesions comprised only 0.2% of the epithelial compartment, they displayed geographic diversity, immune exclusion, metabolic rewiring, and DNA copy number changes among lesions and normal fallopian tube epithelium discovered alterations in STIC-associated genes and the pathways they control. In sum, this platform provides a comprehensive 3D atlas of early neoplastic transformation, yielding mechanistic insights into tumor initiation and informing clinical screening strategies for detecting cancer precursors in whole organs at cellular resolution.

## INTRODUCTION

Ovarian cancer is the most lethal gynecological malignancy, with high grade serous carcinoma (HGSC) accounting for the majority of the cases^1–6^. Accumulating evidence supports the fallopian tube, and serous tubal intraepithelial carcinoma (STIC), as the primary precursor of ovarian cancer^7–16^. This multi-organ progression from the fallopian tube to the ovary is unique among cancers, and its discovery has spurred research into the progression of STICs to invasive HGSC^17–23^. Yet, our current knowledge of ovarian cancer precursors largely stems from studies involving clinical specimens from individuals with ovarian cancer, gynecologic abnormalities, or from high-risk individuals possessing genetic risk factors^24–27^. Consequently, our knowledge of ovarian cancer precursors in wholly non-diseased specimens is limited^28–31^.

STIC is diagnosed incidentally under microscope following the pathological criteria previously reported^32,33^. STIC lesions consist of atypical and multi-layered epithelial cells, with detectable mitotic figures and higher proliferative activity as compared to background epithelium. Alongside STIC, another related lesion emerges as a “p53 signature,” which is defined as a minute stretch of morphologically unremarkable epithelium but harboring *TP53* mutations. The biological and clinical significance of p53 signatures is unclear and whether they represent the precursor lesions of STIC awaits further molecular studies. STICs are more commonly identified following the Sectioning and Extensively Examining the Fimbriated End, or SEE-FIM, protocol that has been adopted as a more thorough way in sampling fallopian tubes in clinical practice^34–36^. However, the diagnosis of STICs is solely based on 2D examination of tissue sections, and consequently, as little as 1% of tubal tissues is microscopically examined by a pathologist as the bulk remains in archived tissue blocks^37–40^. As a result, the actual prevalence of STICs remains unknown and previous studies reported a wide range of STIC incidence, ranging from 11-61% in HGSC patients^41^, 0-11.5% in asymptomatic BRCA1/2 germline mutation carriers^5,37,42,43^, and <1% in individuals without ovarian cancer or genetic risk factors^44–47^. A significantly higher incidence was noted when the tissue blocks were flipped over and additional sections examined^43^, supporting the idea that current sampling lacks the sensitivity to exhaustively detect STIC lesions. Therefore, automated and exhaustive three-dimensional assessments are essential to resolve the spatial distribution and prevalence of rare and microscopic lesions, and to analyze their unique properties as the earliest stage of ovarian tumorigenesis^48–56^.

To address this gap, we developed a novel framework to comprehensively screen entire organ donor fallopian tubes for ovarian cancer precursors. Organ donation for scientific research is a precious resource that provides essential access to tissues unaffected by cancers and other abnormalities generally present in clinical specimens. Donor tissues have emerged as crucial for characterization of precancer frequency and molecular characteristics in organs including pancreas and colon^57,58^. Importantly, because detection of STICs requires consideration of H&E, p53, and Ki67, existing 3D pathology workflows that rely solely on H&E are insufficient to reliably identify these lesions. To overcome this, we developed a pipeline to combine H&E-based cellular morphology with signal intensity from co-registered p53 and Ki67-stained IHC images to automatically highlight hundreds of potential precancerous lesions in a format easily reviewed by expert pathologists and amenable to further integration of multi-omics at regions of interest. This integration allowed an exhaustive and precise 3D mapping of microscopic p53 signatures, proliferative dormant and active STICs in whole human fallopian tubes at cellular resolution.

While previous works have suggested these lesions are rare in low-risk populations^44–46^, using our automated and whole organ-scale workflow we find multiple p53 signatures, proliferative dormant or active STICs in all donor samples analyzed. Digital simulation of the SEE-FIM protocol in these donor organs explains their apparent elusiveness, revealing that the standard SEE-FIM protocol would detect less than half of the precursor lesions found here. To reduce the false-negative rate below 25%, 150-250 equally spaced sections would be required. This is dramatically higher than the 10-20 sections typically analyzed in SEE-FIM and explains the historic lack of evidence of STIC lesions in non-diseased fallopian samples.

Next, we further extended our workflow to integrate spatial proteomics, spatial transcriptomics, and spatial metabolomics to perform deeper molecular profiling specifically in regions within whole donor fallopian tubes that contained precancerous lesions. Using a 25-plex CODEX panel, we found that isolated STICs do not possess a unique immune microenvironment, unlike STICs found in the clinic, suggesting that immune evasion may not be an early hallmark in STIC progression. Using stimulated Raman scattering hyperspectral imaging (SRS-HSI) based spatial metabolomics approach^59^, we identified oxidative stress and increased rigidity that promotes malignant transformation. We also found increased nicotinamide adenine dinucleotide to oxidized flavin adenine dinucleotide (NADH/FAD) ratio in lesion cells compared to surrounding normal epithelial cells, suggesting the lesion epithelia subjects to oxidative stresses and rewires its metabolism towards glycolysis. Finally, integration of Visium spatial transcriptomics revealed significant and spatially confined upregulation of genes essential to cell proliferation, mitotic progression, and chromatin remodeling within the proliferative active STIC epithelium. Lastly, copy number alteration inference in proliferative active STIC showed chromosomal imbalances^60^.

In sum, through integration of high-resolution 3D imaging with molecular profiling, this study reveals the first detailed map of ovarian precancerous lesions in grossly unremarkable fallopian tubes and provides a framework for advancing the understanding of the earliest stages of ovarian cancer development.

## RESULTS

### Construction of cohorts of 3D-microanatomically labelled human fallopian tubes for assessment of ovarian cancer precursors

To assess the presence of ovarian cancer precancerous lesions, including p53 signatures and STICs, in gynecologically healthy women, we developed a pipeline to collect intact donor fallopian tube samples. Samples were procured through the network for pancreatic organ donors (nPOD) from individuals with no documented gynecological disease and no genetic risk factors for ovarian cancer (Fig. 1A). Organs were accepted if donors suffered no abdominal trauma and if warm ischemic time (WIT) was <16 h.

**Fig. 1.**
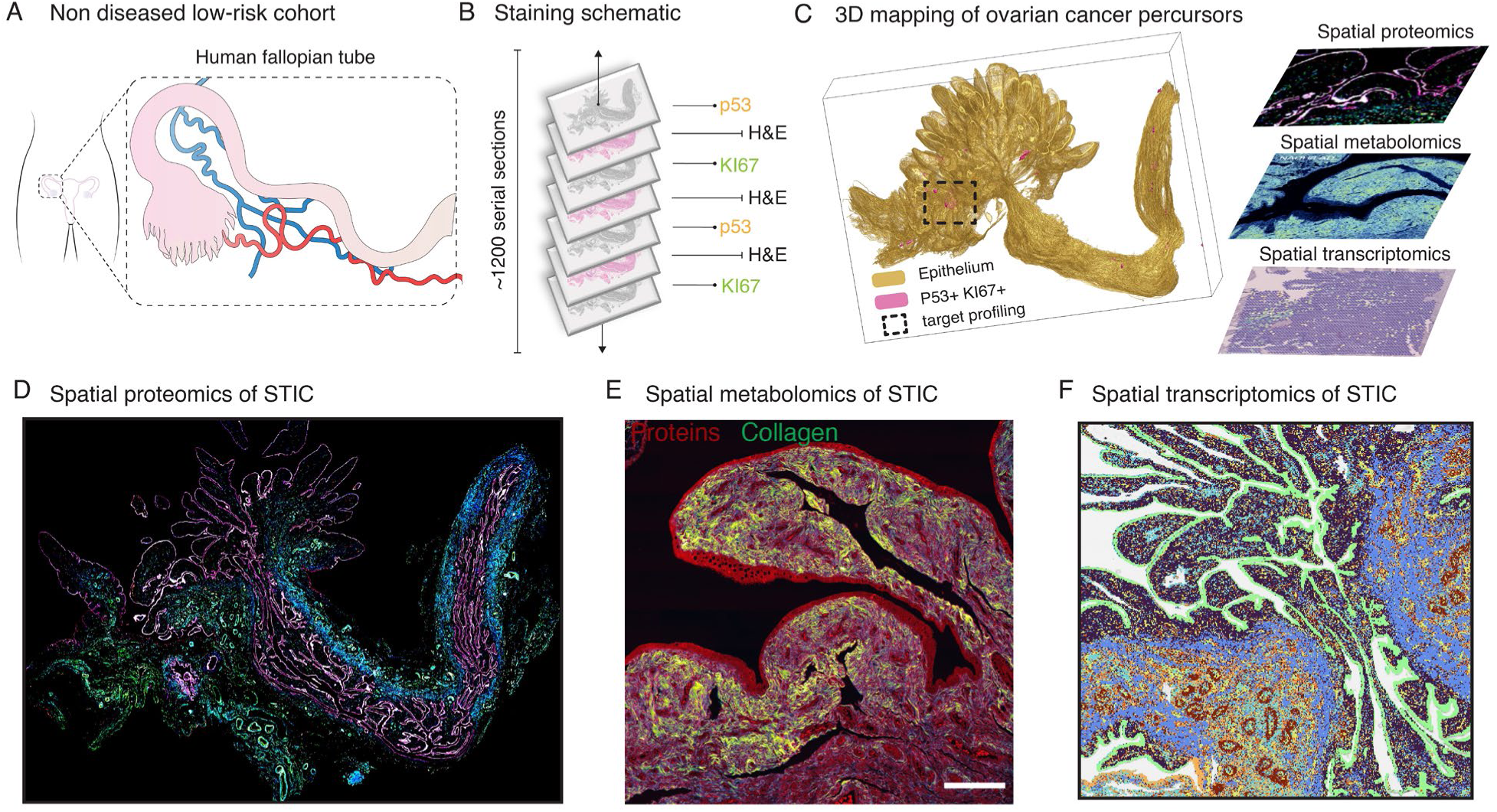
A novel workflow for whole-organ screening of organ donor fallopian tubes for detection and molecular characterization of rare and microscopic lesions. **(a)** Fallopian tubes were obtained from low-risk, non-diseased donors via the nPOD network. **(b)** Specimens were surgically resected, histologically sectioned, stained with H&E and digitized at high resolution. A subset of the sections was stained with *p53* and proliferation marker *Ki67* using IHC. **(c)** H&E- and IHC-stained sections were reconstructed into digital 3D volumes using nonlinear registration. A semantic segmentation algorithm was trained to label tissue components in the H&E images, and starDist was used to segment nuclear boundaries. A supervised algorithm was used to locate 3D regions containing HGSC precursors 3D. **(d-f)** Pathologist-validated lesions were further molecularly profiled using **(d)** spatial transcriptomics, **(e)** stimulated Raman scattering hyperspectral imaging (SRS-HSI) based spatial metabolomics, and **(f)** spatial proteomics.

Fallopian tubes were processed into formalin fixed, paraffin embedded (FFPE) blocks and exhaustively serially sectioned at a thickness of 4 microns. One in every two sections was stained with H&E, one in every eight sections was IHC stained using *p53,* and one in every eight sections was IHC using *Ki67* (Fig. 1B). Stained slides were imaged at 20x resolution (0.5 micron/pixel) using a Hamamatsu S210 scanner, stored as NDPI files, and post-processed into tiff image files. The mean number of sections cut for each human fallopian tube was 981, median 999, maximum 1373, and minimum 601. The average dimensions of the convoluted fallopian tubes in the FFPE blocks were 2.43 cm x 2.26 cm x 0.5 cm, median 2.34 cm x 2.10 cm x 0.5 cm. The average total volume per fallopian tube was 0.68 cm^3^, median 0.75 cm^3^. For context, the median volume of fallopian tube sampled by a single of whole slide is 0.00075 cm^3^ (=0.75cm^3^/999). The mean and median number of cells per fallopian tube was 438.3 million and 509 million, respectively. To preserve DNA, RNA, and proteins, unstained sections were mounted on plus slides and stored with desiccant packets at -20°C.

We trained three deep learning models to semantically segment the fallopian tube microanatomy. The first model segmented eight structures from the H&E-stained images: tubal epithelium, mesothelium, blood vessels, stroma, fat, nerve, rete ovarii, and background. The second model sub-classified the fallopian tubal epithelium into secretory and ciliated epithelial cells. The third model masked locations of positive *p53* and *Ki67* signals on the IHC images. Alignment of the H&E and IHC segmented images into a volume via nonlinear image registration^55,61^ enabled automatic identification of secretory epithelial cells featuring *p53+*/*Ki67*+ and *p53+*/*Ki67*-signal. p53 staining positivity was defined herein as the staining pattern consistent with a *TP53* missense mutation using the criteria previously reported^62^. *Ki67* positivity was defined as the *Ki67* labeling index was significantly higher than that of the adjacent or background epithelium. At these regions, we exported stacks of high-resolution registered 2D images, allowing human validation of detected lesions. A total of 1,285 deep learning-highlighted *p53+*/*Ki67*+ and *p53+*/*Ki67*-epithelium locations, with a mean of 257 and a median of 211 per fallopian tube, were automatically detected by our algorithm and then manually validated by pathologist experts. Highlighted locations were categorized as proliferative active STICs, proliferative dormant STICs, p53 signatures, or non-lesions (Fig. 1C).

Following detection of epithelial lesions, intervening unstained slides were used for deeper profiling. To understand the immune microenvironment of the proliferative active STIC, we applied a CODEX panel of 25 antibodies for WSI proteomics analysis (Fig. 1E). To understand the metabolic changes, we used spatial single-cell metabolomics (Fig. 1E). Lastly, to study gene expression variations and infer copy number alterations, we applied 10x Genomics Visium Cytassist (Fig. 1F).

### 3D characterization of the microanatomy of the human fallopian tube and STIC lesions

To comprehensively study the microanatomy of the fallopian tube in organ donor samples, we analyzed the results of the registered, segmented H&E images (Fig. 2A). High-grade serous tumors primarily originate from secretory epithelial cells in the human fallopian tube^63^, highlighting the importance of understanding the composition and spatial arrangement of secretory epithelial cells in pre-and post-menopausal non-diseased fallopian tubes. Here, we analyzed 175.1 million pre-menopausal epithelial cells and 112.3 million post-menopausal epithelial cells. We produced z-projection heatmaps and 3D reconstructions, conveying the marked convolutions of the fallopian tube epithelial and the intermixing of secretory and ciliated epithelial cells (Fig. 2B and 2C). We found on average higher composition of secretory epithelial cells in post-menopausal (76% secretory, 24% ciliated) women compared to pre-menopausal women (58% secretory, 42% ciliated (Fig 2D).

**Fig. 2.**
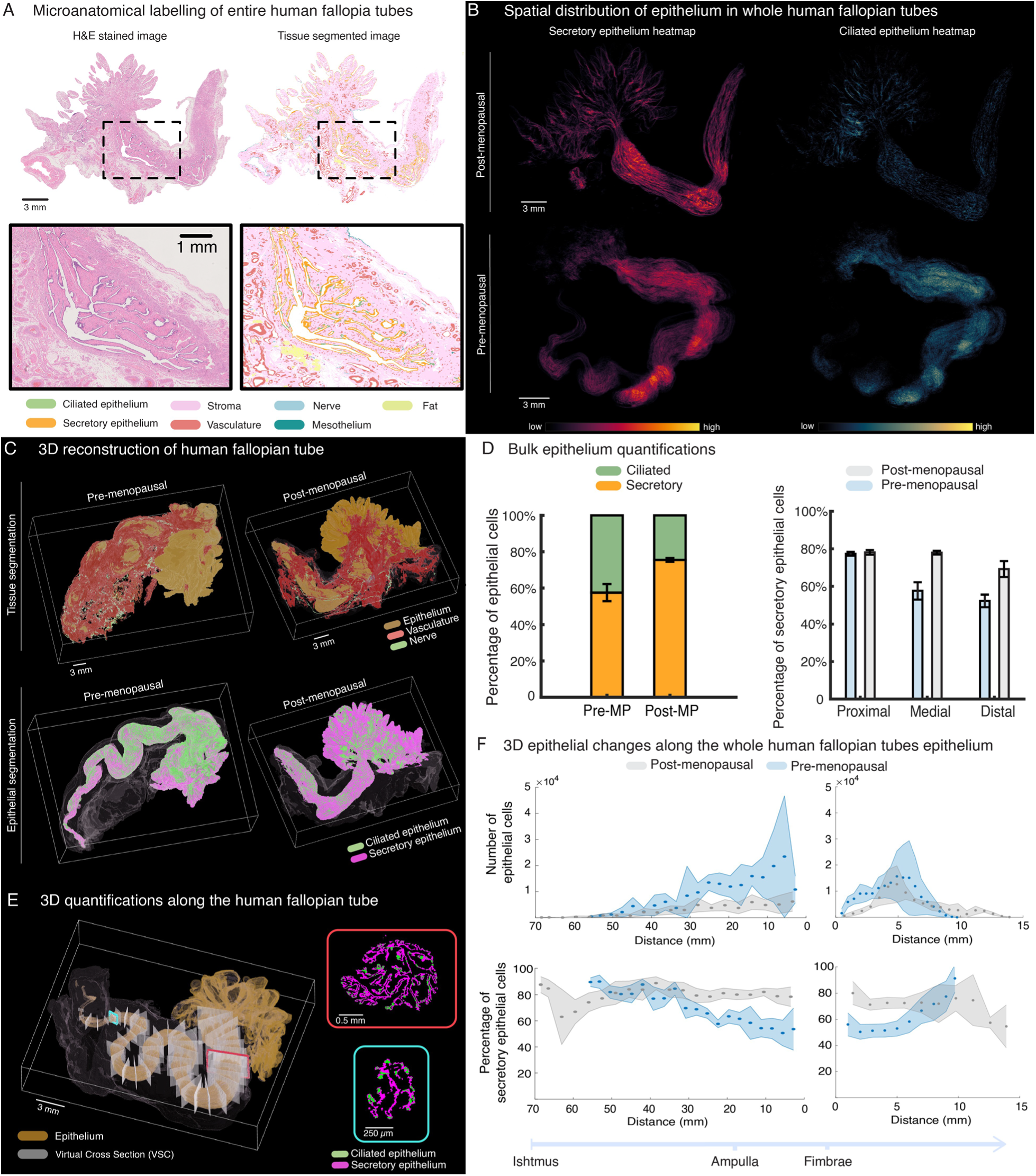
3D analysis of tissue and cellular components of the human fallopian tube. **(a)** The CODA semantic segmentation platform was used to label microanatomical components of the human fallopian tubes from H&E-stained images, including secretory epithelium, ciliated epithelium, mesothelium, blood vessels, stroma, fat, and nerve. **(b)** 2D heatmaps obtained form best projections of the whole stacks of labeled images showing the differences in the human fallopian tube’s epithelium, whereby post-menopausal samples showed less ciliated epithelium than pre-menopausal samples. **(c)** Major tissue components of the fallopian tube reconstructed using the 3D CODA mapping platform, which revealed high vascularization of human fallopian tubes (top panels). 3D rendering of epithelial subtypes confirmed the presence of higher secretory epithelial populations in post-menopausal samples when compared to pre-menopausal samples (bottom panel). **(d)** Bulk tissue composition plots of ciliated and secretory epithelium in post- and pre-menopausal human fallopian tubes (top panel), and plotted compared to proximal, medial, and distal locations to ovaries (bottom panel). **(e)** Center path of each human fallopian tube was computed to measure variance in epithelium across each specimen. **(f)** Cross sectional analysis of the fallopian tubes revealed an increase in secretory epithelial cells along the entirety of the post-menopausal fallopian tubes, when compared to pre-menopausal cohort.

Our 3D maps of whole fallopian tubes allowed us to computationally generate “virtual” sections of selected orientation (e.g. orthogonal to the main axis of the fallopian tube). To generate virtual sections along the length of the fallopian tube, we skeletonized each specimen by calculating the center path along the convoluted tubal lumen. At each cross section along the tube, we calculated the distance to the ovary (defined at the tip of the fimbriated end), and categorized this distance as proximal, medial, or distal. We visualized (Fig 2C, bottom) and quantified (Fig 2E, right) the distribution of secretory cells to show that the drop in ciliated cell content from pre- to post-menopausal primarily affects the locations on the fallopian tube medial and distal to the ovary, with similar composition of ciliated cells proximal to the ovary across age groups (Fig. 2E, right). We further quantified the secretory and ciliated cell composition as a function of distance along the center path, for precise sampling by generating thousands of orthogonal virtual cross sections along the epithelium (up to 10,255 virtual sections per sample). For each cross section, we quantified the overall frequency of secretory and ciliated epithelial cells from the distal isthmus to the proximal fimbriated end.

As the majority of STICs originate from secretory epithelial cells^64^, understanding their spatial distribution and age-associated changes is critical for understanding early ovarian tumorigenesis. Using our workflow to quantify the normal epithelial composition 3D, the data revealed that, in post-menopausal tubes, the proportion of secretory cells increases sharply toward the fimbriated end. In contrast, pre-menopausal tubes demonstrate a distal decrease in secretory cell percentage with a concomitant increase in total epithelial cell number due to expansion of ciliated cells. These data indicate that menopausal status substantially remodels the cellular composition of the distal tube towards a more secretory epithelial cell landscape, potentially influencing the local risk for neoplastic precursor lesions.

### 3D mapping of lesions in the non-diseased human fallopian tube epithelium

To implement a strategy for detailed 3D mapping of epithelial lesions in average-risk, nondiseased human fallopian tubes, samples were alternately stained with H&E, *p53* IHC and *Ki67* IHC (Fig. 3A). Implementation of segmentation of the H&E and IHC images allowed automated detection of p53 signatures, proliferative dormant STICs, and active STICs following standard clinical definitions (Fig. 3B). 3D volumetric renderings of these lesions convey their microscopic size and wide range of 3D morphology (Fig. 3C).

**Fig. 3.**
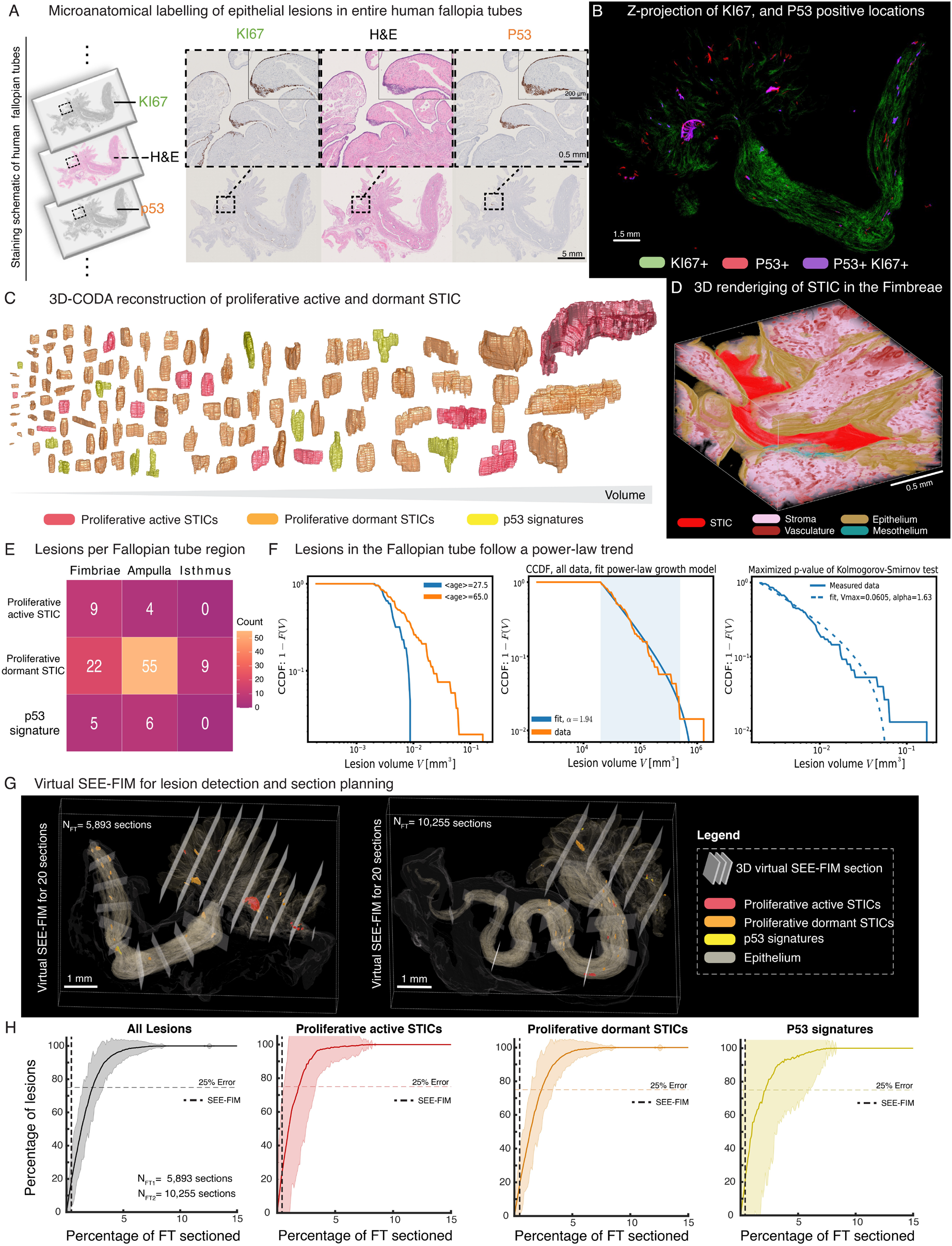
3D mapping of ovarian cancer precursors reveals high lesion burden with spatially patterned, scale-free growth. **(a)** Integrated pipeline combining Immunohistochemistry staining (*Ki67* and *p53*) and 3D computational reconstruction to map lesions across entire fallopian tubes (n=5). **(b)** Z-projection demonstrating detection of proliferative *Ki67*+ and *p53*+ aberrant foci. **(c)** Pathologist-expert validated lesions were labelled as proliferative active STIC, proliferative dormant STICs, and p53-signatures, and 3D rendered. **(d)** Volumetric rendering of a proliferative active STIC within fimbriae architecture (scale: mm). **(e)** Spatial distribution of lesions, showing high lesion burden and sparse distribution of lesions in the human fallopian tubes**. (f)** Complementary cumulative distribution functions (CCDFs) of lesion size follow a power-law trend across all tube regions. **(g)** Example of 3D virtual SEE-FIM computed for a post-menopausal sample containing 5,893 virtual sections. **(h)** 3D virtual SEE-FIM procedure was computed for incrementally increasing number of virtual sections. Percentage of detected combined lesions, proliferative active STICs, proliferative dormant STICs, and p53 signatures was calculated for each equally distant virtual section count. The number of sections used in standard SEE-FIM procedures is indicated by the black or orange vertical lines.

We identified and 3D mapped 99 STICs, including 13 proliferatively active STICs, 86 proliferative dormant STICs, and 11 *p53* signatures across 5 nondiseased whole human fallopian tubes from 5 distinct donors (Fig. 3D). According to menopausal status, we observed an average of zero STICs, 8.5 proliferative dormant STICs, 3 p53 signatures in pre-menopausal samples (Table S1, S2, S3, and S4). Our data revealed that ovarian precancerous lesions were present in 80% of the examined fallopian tubes (Tabel S2, S6). In post-menopausal samples, we observed an average of 4.33 STICs, 23 proliferative dormant STICs, 1.67 p53 signatures. Notably, one post-menopausal sample contained an unusual high number of lesions: 5 STICs, 37 proliferative dormant STICs, and 4 p53 signatures. The most common lesion we identified proliferative dormant STIC, and the most common location found to contain proliferative dormant STICs was the ampulla (55 lesions), compared to the fimbriated end (22 lesions) and isthmus (9 lesions). The most common location to contain proliferative active STICs was the fimbriated end (9 lesions) followed by the ampulla (4 lesions) and no active STICs were found in the Isthmus. We found p53 signatures in both the fimbriated end (5 lesions) and the ampulla (6 lesions).

The occurrence of STIC, proliferative dormant STIC, and p53 signature lesions was higher in post-menopausal donors compared to pre-menopausal donors. STICs were detected in 67% of post-menopausal donors but were absent in pre-menopausal cases. proliferative dormant STICs were detected in 100% of post-menopausal donors and in 50% of pre-menopausal donors (Table S5). The p53 signature was present in 67% of post-menopausal and 50% of pre-menopausal donors. When combining all donors, the overall prevalence was 40% for STIC, 80% for proliferative dormant STIC, and 60% for p53 signature.

### Growth model of epithelial lesions in average-risk intact human fallopian tube samples

Our previous work in mathematical modelling of precancerous lesions of the human pancreas (PanINs) suggested that a simple growth law allowing each anatomically separate lesion to grow at a constant rate is insufficient to explain the very large lesions found in our cohort^65^. Explaining the size distribution required additional actions such as lesion splitting and lesion merging, which was confirmed by genomic data and suggested that some large PanINs are composed of multiple clones that collided within the pancreatic ductal system^66^. In contrast, simple growth laws explain the lesion distribution in healthy fallopian tubes, for which a maximized p-value using of Kolmogorov-Smirnov test resulted in a V_max_ of 0.0605 mm^3^ and an exponent of α=1.63 (p = 0.57, Fig. 3F, right panel). This result suggests, unlike PanINs, a lack of polyclonality in the microscopic lesions found in this cohort of organ donor fallopian tube samples (Fig. 3F, middle panel).

### Development of virtual SEE-FIM for statistical determination of fallopian tube sampling guidelines

We asked why previous SEE-FIM-based assessments have not detected this high occurrence of lesions. To quantify the impact of subsampling when detecting ovarian cancer precursors, we virtually implemented a virtual SEE-FIM protocol. We generated longitudinal sections at the fimbriated end and transverse sections along the remainder of the ampulla and isthmus, as done in the clinic. To show the ability of SEE-FIM to identify lesions, including proliferative active STICs, dormant STICS, and p53 signatures were highlighted in red, orange and yellow, respectively (Fig. 3G, and Fig. S2D).

First, we simulated current SEE-FIM guidelines via collection of 20 equally spaced virtual sections, representing approximately 0.25% volume of the entire organ. Within the extracted sections, SEE-FIM was able to identify 10.8% of all lesions, 14.6% of STICs, 10.1% of proliferative dormant STICs, and 11.9% of p53 signatures. These results reveal that conventional SEE-FIM protocol may significantly underestimate the true incidence of precursor lesions in human fallopian tubes. We determined that approximately 2.3%, or 186 tissue sections, of the whole fallopian tube would need to be assessed to accurately identify all lesions with <25% error (Fig. 3H, left panel, Fig. S2E, top left panel). Splitting by lesion type, we determined that to accurately identify STICs, proliferative dormant STICs, and p53 signatures with 25% error, 1.8% (149 sections), 2.4% (190 sections), and 2.2% (174 sections) of the fallopian tube would need to be assessed, respectively (Fig. 3H, Fig. S2E).

### Spatial protein marker profiling of STIC in non-diseased human fallopian tubes

To study the microenvironment surrounding the proliferative active STIC identified in 3D, we applied a panel of 25 protein markers using CODEX multiplexed imaging. We applied nucleus and cell body segmentation to identify 972,276 cells across the whole slide image (Fig.4A-B)^67^. We performed unsupervised clustering to obtain 30 distinct clusters, which we annotated and combined the clusters into 19 relevant cell phenotypes using previously established methods (Fig. 4C)^68–70^. These cellular phenotypes included STIC, epithelial cells, immune cell phenotypes (T cells, B cells, macrophages, neutrophils, dendritic cells), stromal cells (fibroblasts, smooth muscle cells), and tumor associated macrophages (TAMs), shown spatially in Fig. 4D. The protein expression matrix (Fig. 4E) and protein markers interactions^71^ (Fig. 4F) illustrate that epithelial and proliferating epithelial cells interact with EpCAM, Pan-CK, and *Ki67*. We labelled activated and memory T cells by CD3, CD4, CD8, and CD45RO, with additional links to IFNG and CD44. We identified B cells via CD20, and dendritic or APC populations by HLA-DR, CD11c, and CD141. Macrophages (TAMs) and monocytes associate with CD68 and IDO1, neutrophils with MPO, and endothelial cells with CD31. Mesenchymal and myofibroblast identities are confirmed by Vimentin and SMA. These specific interactions validate the correct phenotypic annotation in the dataset.

**Fig. 4.**
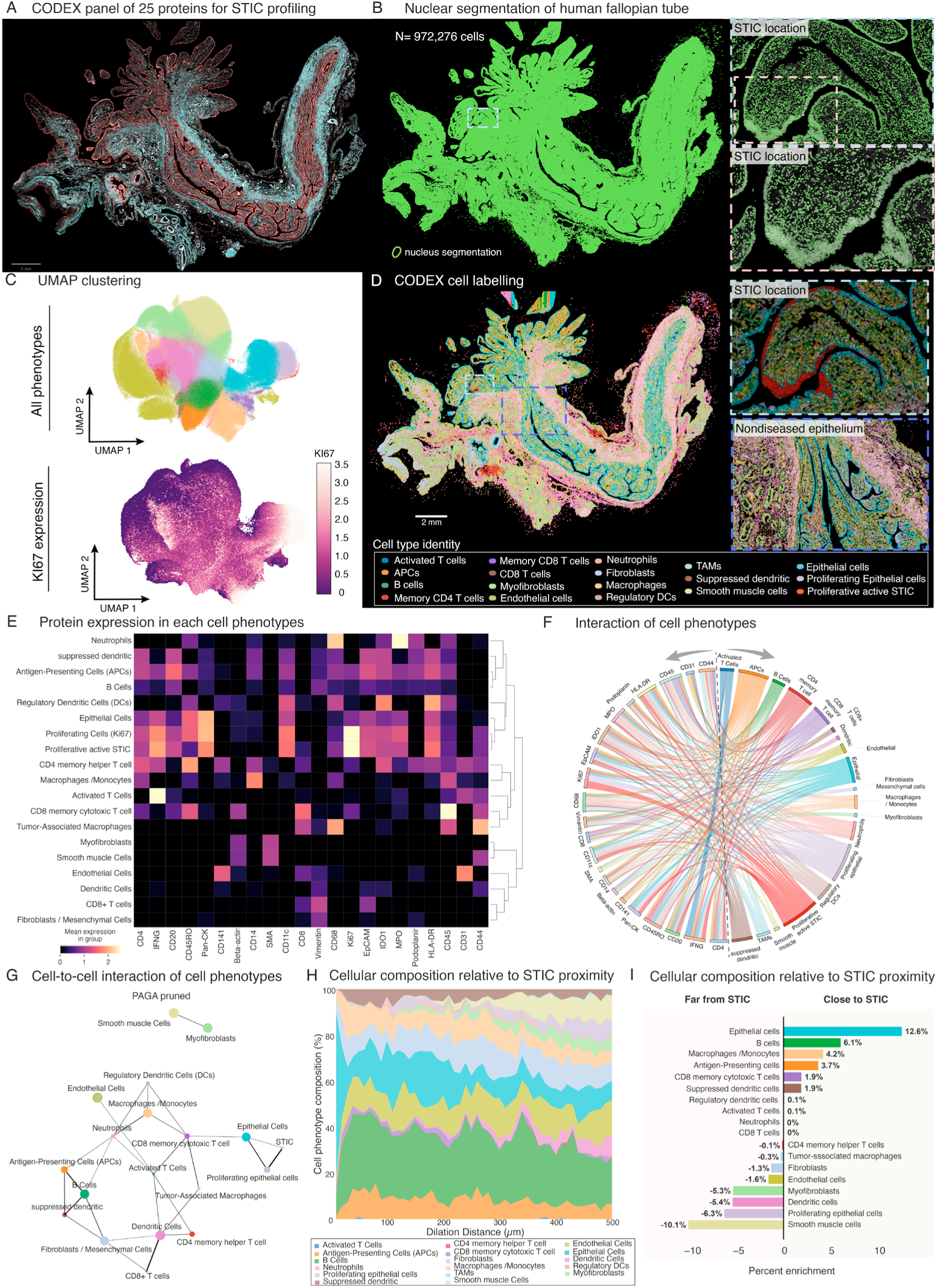
Proliferative active STIC lesion creates an immunosuppressive niche through altered cellular crosstalk and spatial reorganization of the tumor microenvironment. **(a)** Multiplexed CODEX imaging reveals protein expression patterns across 25 markers in proliferative active STIC. **(b)** Single-cell resolution mapping of fallopian tube epithelium (972,276 nuclei). **(c)** UMAP clustering identifies distinct cellular phenotypes, with proliferative (*Ki67*+) epithelial-immune clusters enriched in STIC. **(d)** Cell-type-specific protein signatures highlight metabolic and immune checkpoint dysregulation in STIC and adjacent stroma. **(e)** PAGA network analysis uncovers rewired interactions between stromal fibroblasts and immunosuppressive myeloid populations in STIC. **(f)** Spatial profiling demonstrates immune exclusion, with cytotoxic T cells displaced and suppressive dendritic cells recruited near STIC. **(g)** Quantification of immune-stromal shifts across increasing distances from STIC core. **(h-i)** Comparative cellular landscapes reveal modest immune localization near the STIC.

Partition-based Graph Abstraction (PAGA) of single-cell proteomics in Fig. 4G^72^ showed interactions between the distinct cell phenotypes. STIC cells were closely associated with proliferating epithelial cells, supporting a trajectory consistent with malignant epithelial progression, while showing no direct connectivity to any other cell populations. Analysis of the PAGA connectivity map further identified interconnected immune cell populations comprising tumor-associated macrophages (TAMs), regulatory dendritic cells (DCs), activated T cells, and CD8+ memory cytotoxic T cells, suggesting potential immune coordination mechanisms.

PAGA analysis identified an immune network connecting TAMs, regulatory DCs, and activated T cells to CD8+ memory cytotoxic T cells. Additionally, CD8+ T cell connection to epithelial cells further bridging to STIC populations. This CD8+ T cell population’s dual connectivity to both immunosuppressive cells and epithelial cells may indicate potential compromised immune surveillance^73–75^. While these topological relationships require functional validation, their organization suggests structurally relevant cellular interactions potentially governing STIC maintenance^76^. This immunological landscape closely mirrors established observations in ovarian cancer literature, where TAM enrichment and regulatory immune cell infiltration consistently correlate with tumor progression and poor clinical outcomes^77–80^.

To further validate the results obtained from the PAGA graphs, spatially resolved protein profiling was implemented to assess immunosuppressive expression within these interacting cell populations. Thus, to spatially assess the STIC microenvironment, STIC mask was generated and consequently dilated in 10-micron increments up to 500 microns from the STIC boundaries (Fig. S3B). For each distance interval computed, cellular composition was estimated and visualized (Fig. 4H). To quantify the differences in cellular composition relative to proximity to STIC location, comparison between regions close to the STIC (less than 100 microns) and regions distant to the STIC (between 100 and 500 microns) was performed. Proximity to the STIC showed modest enrichment in macrophages or monocytes, B cells, antigen-presenting cells (APCs), suppressed dendritic cells, and CD8 memory T cells (Fig. 4I).

Spatial CODEX analysis identified an immunoregulatory microenvironment surrounding STIC lesions, with an increase of macrophages, regulatory dendritic cells, and CD8 memory T populations in proximal regions (Fig 4I). This validated the PAGA analysis, which showed direct connectivity between these immune populations and epithelial cell states. The spatial organization of these macrophage and CD8 T-cell populations aligns with the immune modulation observed in early high grade serous ovarian cancer^77^. These findings define the STIC microenvironment as a site of coordinated immune epithelial interactions that may facilitate early lesion persistence. Our results were consistent with those previously published^81^.

### Spatial metabolomics profiling of STIC in non-diseased human fallopian tubes

Existing single cell and spatial transcriptomics data analysis has shown association of fallopian tube epithelium to genes and pathways associated with metabolic regulations in various contexts ^82–84^. With this in mind, we carefully evaluated unsaturated lipid level and redox ratio of preneoplastic epithelial lesions by multimodal two-photon stimulated Raman scattering imaging (SRS) (Figures 1A-C, S1A and S2A-C). We found that the unsaturated lipid level was reduced while the redox ratio indicated by NADH/FAD was increased in the lesion cells compared to the healthy surrounding cells (Figure 1C), suggesting the lesion epithelia subjects to ROS stress and metabolic remodeling towards glycolysis ^85^. Worth noting that the scattered distribution of NADH and FAD signals in lesions (Figure 1C in cyan and magenta) may indicate the fragmented mitochondria with compromised metabolic function. We further applied hyperspectral SRS imaging and lipid subtype detection^59^ to perform an in situ lipidomic analysis and identified that lesions represented a distinct lipid profile manifesting in upregulated ceramide/PE and PC/PE ratio (Figures 1D-F and S1B), aligning with a study showing the disturbed homeostasis of ceramides and phospholipids in abnormal epithelial context ^86^.

Interestingly, Raman spectra showed distinct changes of lipid profile in different types of lesions (Figures S2D-F), suggesting the lipid metabolism is highly sensitive to the lesions and different lipid profile may represent the trajectory of lesion development. Altogether, our results provide new insights into the molecular mechanisms underlying the lesions of fallopian tube epithelium.

### Spatial transcriptomic profiling of STIC in nondiseased human fallopian tubes

Using the CODA IHC-based deep learning method, we profiled STIC with spatial transcriptomics (Fig. 6A). STIC location was processed using Visium Cytassist for whole transcriptome profiling. Curation of the spatial spots identified STIC epithelial spots in red and non-STIC epithelial spots in green (Fig. 6B).

Differential gene expressions of the proliferative active STIC against normal adjacent epithelium were obtained and shown in volcano plot (Fig. 6C). The upregulated genes in STIC, including *KIF1A*, *TUBB2B*, *DLGAP5*, *BUB1*, *KIF2C*, *CDCA8*, *CDC20*, *CCNF*, *CCNB1*, and *PBK*, suggest dysregulated cell cycle progression, mitotic spindle function, and chromosomal instability, which are seen in high-grade serous ovarian cancer^87–97^. Immune-related genes like *ULBP3* and *BTNL2* may contribute to immune evasion, while *JUN* and *NOX4* could promote survival and oxidative stress responses^98–101^. The presence of *HNF4A*, *TFAP2A*, and *ADAM12* further supports a link to ovarian carcinogenesis through transcriptional deregulation, cellular differentiation, and extracellular matrix remodeling^102–104^. These findings reinforce STIC’s role as a precursor to high-grade serous carcinoma, with key drivers of malignancy already active^105^. Comparative analysis revealed significant upregulation of genes such as *GPX2* (implicated in oxidative stress response) and *HIST1H1D* (a chromatin regulator) in STICs (Fig. 6D), mirroring patterns observed in advanced ovarian tumors^106,107^.

Pathway analysis using the Hallmarks gene sets^108^, and the suppressed and activated pathways were computed (Fig. 6E-F). Hallmark pathway analysis revealed enrichment of several cancer-associated pathways in STICs, including spermatogenesis, *G2M* checkpoint, *KRAS* signaling, *E2F* targets, oxidative phosphorylation, and *TNFα* signaling via *NFκB*^109–116^. These activated pathways are consistent with clinical observations of early oncogenic signaling in STIC lesions that precede invasive high-grade serous carcinoma development^117^. Gene set enrichment analysis profiles confirmed significant enrichment of proliferation-associated pathways and G2M checkpoint genes (Fig. 6G), showing dysregulated cell cycle characteristics of both STICs and invasive ovarian cancers.

To investigate chromosomal instability in proliferative active STIC, copy number analysis (CNA) was inferred from the spatial transcriptomics data (Fig. 6H-I, Fig. S6F), which revealed gains in chr6p22, chr6p21, chr1p32, and chr16p13, and losses in chr17p13, chr9q33, chr9q34, chr22q11, chr22q12, and chr22q13. These results align with clinical genomic studies showing that copy number alterations and genomic instability are early events in STIC lesions^15,109,118,119^. Chromosomal 6 gains and chromosomal 22 depletions were spatially located on the STIC (Fig. 6K). Notably, gains in chr6p, which harbors immune-related genes, have been linked to immune evasion and tumor progression in ovarian cancer, while losses in chr17p13, encompassing *TP53* and are associated with impaired DNA damage response and genomic instability^119^. These alterations may collectively contribute to early malignant transformation and aggressive phenotypes in STIC lesions.

To further explore chromosomal alterations, we also applied inferCNV (Fig. S6F) and identified chromosomal gains in chromosomes 1, 6, 8, 16, and 19. Chromosomal losses were detected in chromosomes 4, 9, 13, 15, 17, 18, and 22. Comparison to a large cohort study of 47 patients with proliferative active STICs^109^, which showed chromosomal gains in chromosomes 1, 2, 3, 6, 7, 8, 10, 12, 16, 19, and 20; and chromosomal depletions in chromosomes 4, 5, 6, 7, 8, 9, 11, 13, 15, 16, 17, and 22. Similarly, genes altered in these regions include *TP53* (chr17p13), *MYC* (chr8q24.21), *CCNE1* (chr19q12), *CDKN2A/CDKN2B* (chr9p21), *BRCA1* (chr17q21), and *NF2/TIMP3* (chr22q12-13).These chromosomal targets highlight pathways associated with cell cycle regulation, DNA repair, and immune modulation. Conversely, the large patient cohort study also reported unique gains in chr2, chr3, chr7, chr10, chr12, and chr20, and unique losses in chr5, chr7, chr8, and chr11, not observed in our nondiseased, average-risk donor cohort analysis. These regions encompass genes such as PIK3CA/MECOM (chr3q26), ETV6/FOXM1 (chr12p13), and APC (chr5q22), which are involved in PI3K signaling, transcriptional regulation, and tumor suppressor pathways.

## DISCUSSION

Here, we present a workflow for quantitative whole organ screening of precancerous lesions combining image registration of multi-plex pathology slides with deep learning-based automated detection of suspicious regions. This workflow enabled us to analyze ∼10,000 whole slide images to identify 99 STICs and 11 p53 signatures across 5 donor samples, as confirmed by a gynecologic pathologist using standard review criteria, demonstrating the importance of quantitative 3D imaging to improve the throughput of pathology review. To the best of our knowledge, the current study shows the first 3D mapping of a whole human fallopian tube at single cell resolution, capable of comprehensively identifying the precancerous lesions and integrate spatial multi-omic profiling at specific regions of interest.

Spatial proteomics uncovered an immune-excluded microenvironment surrounding STIC lesions, with reorganization of stromal and myeloid populations. This immunosuppressive niche mirrors patterns observed in invasive HGSC, supporting the concept that immune remodeling begins during early transformation. Cell–cell interaction analysis revealed that tumor-associated macrophages and regulatory dendritic cells closely interact with proliferative epithelial compartments, potentially facilitating immune tolerance. A recent report also demonstrates immune cold microenvironment associated with the majority of non-BRCA1/2 STICs, further confirming this observation^109^.

Spatial metabolomics revealed higher levels of ceramides in the lesions. Increased ceramide levels have been associated with apoptotic cell death in both homeostatic systems as well as pathological settings as a result of cellular insults including oxidative stress, chemotherapeutic agents, ischemia and radiation^120^. Together with the morphology and redox ratio changes in mitochondria we found, it is possible that ceramide act on mitochondrial pathways to shape the cellular metabolic activity in lesion cells. Actually, ceramide is able to induce apoptosis by recruitment of death receptors to lipid rafts and assembly of channels in the outer membrane of the mitochondria promoting the release of cytochrome^120^. However, further studies are warranted to determine the direct or indirect effects exerted by elevated ceramides in regulating cell metabolism and apoptosis in lesions.

Spatial transcriptomics profiling of the proliferative active STIC revealed alterations observed in STICs, and our results on those 5 specimens were also observed in a larger STIC cohort^109^. In particular, differential gene expression highlighted dysregulated cell cycle, chromosomal instability, oxidative stress, and immune evasion. Pathway analysis showed enrichment of cancer-associated pathways such as G2M checkpoint and KRAS signaling. Copy number alteration analysis of the nondiseased average-risk proliferative active STIC and subsequent comparison to large clinical patient cohort identified overlapping chromosomal alterations with clinical STICs^109^, including chromosomal gains in chromosomes 1, 6, 8, 16, and 19, and losses in chromosomes 4, 9, 13, 15, 17, and 22, which are linked to TP53, MYC, CCNE1, CDKN2A/CDKN2B, BRCA1, and NF2/TIMP3. These chromosomal targets highlight pathways associated with cell cycle regulation, DNA repair, and immune modulation. Chromosomal alteration differences from both cohorts included unique gains in clinical STICs (chromosomes 2, 3, 7, 10, 12, and 20) and losses (chromosomes 5, 7, 8, and 11), which encompass genes such as PIK3CA, FOXM1 and APC, which are involved in PI3K signaling, transcriptional regulation, and tumor suppressor pathways. Nondiseased low-risk STIC displayed a less extensive genomic profile, suggesting distinct molecular landscapes that may reflect differences in progression stages.

Limitations of this study include the modest sample size, which limits precise estimation of lesion prevalence in the general population. This constraint stems largely from the rarity of donor fallopian tube specimens available for analysis, as the samples were obtained from an organ donor network from women at low risk of ovarian cancer. Nonetheless, detection of lesions across all samples highlights the potential for the technical platform reported here to improve current diagnostic protocols, which may substantially underreport the burden of early neoplasia in the fallopian tube. With modifications to the grossing protocol, this approach could be applied to remnant surgical specimens following clinical assessment of 2D sections necessary for patient diagnosis.

This study establishes a scalable framework for 3D mapping of precancerous lesions in whole human fallopian tubes, allowing the concurrent profiling of the lesions using spatial proteomics, metabolomic and transcriptomic technologies. These 3D maps can be used to perform pseudo-time modelling of precancerous lesions and for development of biomaterials that molecularly, functionally, and architecturally resemble human fallopian tubes and their precancerous lesions^53,121^. Future work should expand cohort size, integrate longitudinal samples, and explore how this new knowledge can be translated into clinically meaningful data for future development of effective tools for early diagnosis and strategy for ovarian cancer prevention.

---

**Fig. S1.**
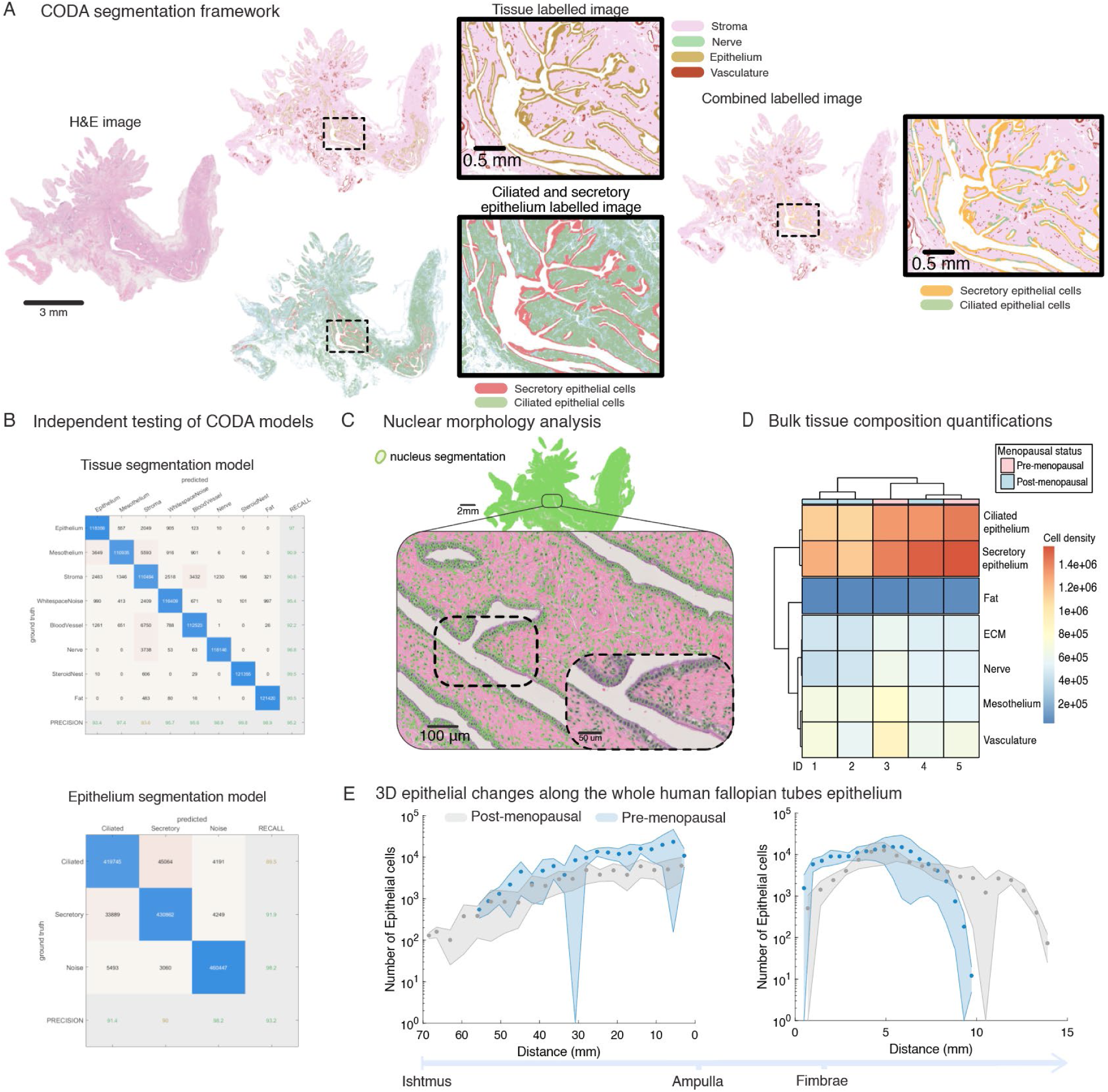
3D-CODA single cell resolution framework to map epithelial cell subtypes. **(a)** Representation of H&E-stained human fallopian tube whole-slide image (left panel). CODA segmented tissues, such as epithelium, mesothelium, blood vessels, fat, nerves (middle panel, top). CODA subtyped the epithelium into secretory epithelium and ciliated epithelium (middle panel, bottom). Combination of the two segmentation models allowed for detailed tissue mapping of whole fallopian tube H&E-stained images (right panel).**(b)** Testing of the segmentation models was performed on independent images from the training dataset. Tissue model showed overall accuracy of 95.2% (top panel) and epithelial subtyping model showed an overall accuracy of 93.2% (bottom panel).**(c)** Nuclear segmentation model was applied to all H&E-stained images of each human fallopian tube to obtain cellular resolved data. **(d)** Bulk cell density was measured for each tissue type of each human fallopian tube. Ciliated and secretory cell density was lower for ID 1 and 2, which contained the most lesions. **(e)** Cross sectional analysis of the fallopian tubes revealed increased in secretory epithelial cells along the entirety of the post-menopausal fallopian tubes, when compared to pre-menopausal cohort.

**Fig. S2.**
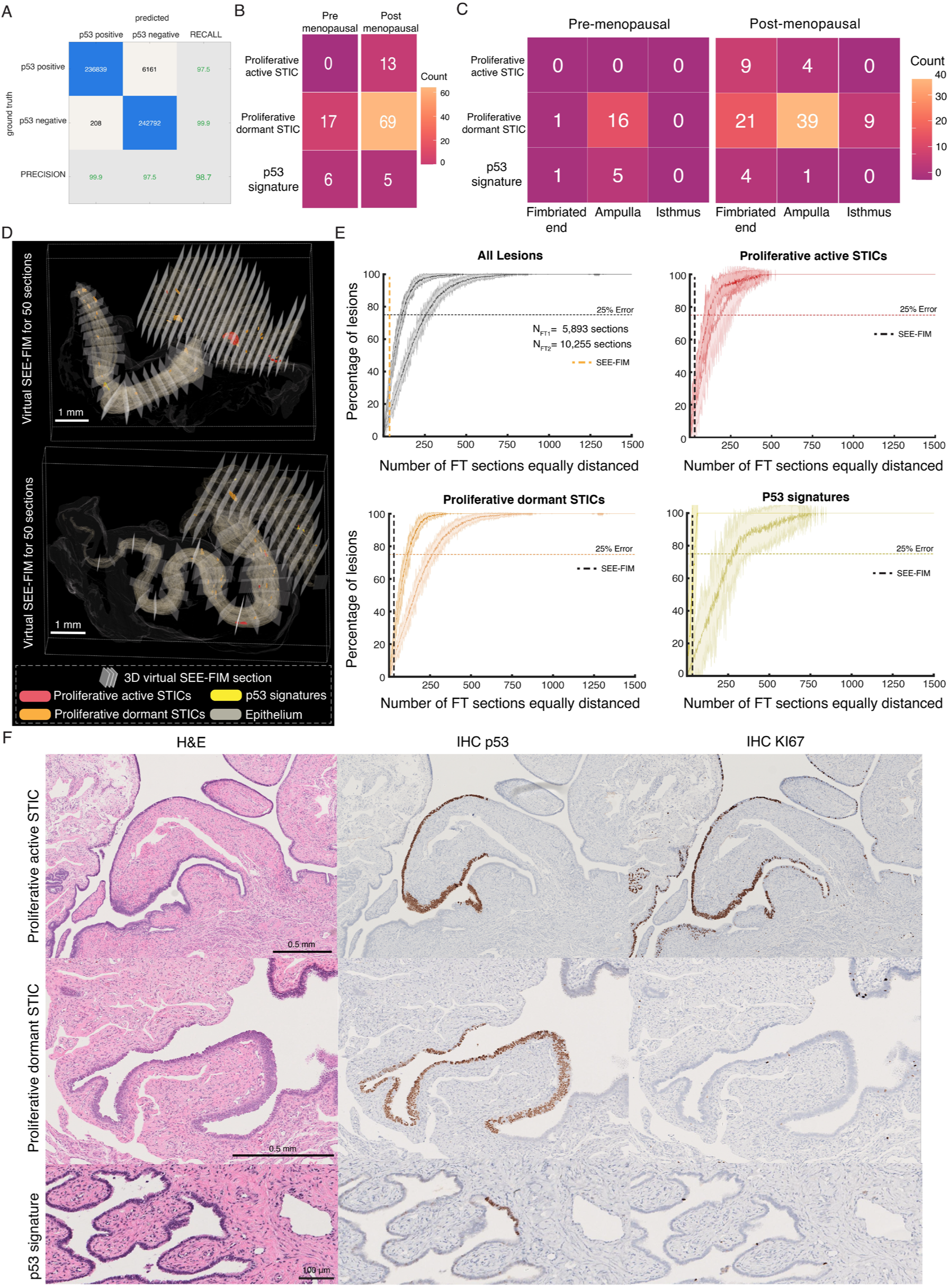
3D spatial and bulk mapping of ovarian cancer precursors. **(a)** CODA segmentation model to annotate IHC positive signal in whole slide images was tested on an independent testing dataset and achieved an overall accuracy of 98.7%. **(b)** Semi-automated method of lesion detection in whole human fallopian tubes showed 13 STICs, 64 proliferative dormant STICs, and five p53 signatures across 3 samples of the post-menopausal cohort. On the 2 samples of the pre-menopausal cohort, we identified zero STICs, 17 proliferative dormant STICs, and six p53 signatures. **(c)** STICs, proliferative dormant STICs, and p53 signatures were separated according to their spatial locations in the fallopian tube. STICs were found only in Post-menopausal samples, with 9 STICs in the Fimbriated end and four STICs in the Ampulla. The majority of proliferative dormant STICs were found in the Ampulla in both pre- and post-menopausal samples, with 16 and 39 proliferative dormant STICS, respectively. The p53 signatures were identified mostly in the Ampulla and Fimbriated end. **(d)** Example of 3D virtual SEE-FIM computed for a post-menopausal sample containing 10,255 virtual sections. **(e)** 3D virtual SEE-FIM procedure was computed for incrementally increasing number of virtual sections. Percentage of detected combined lesions, STICs, proliferative dormant STICs, and p53 signatures was calculated for each equally distant virtual section count. The number of sections used in standard SEE-FIM procedures is indicated by the black or orange vertical lines. **(f)** Example of p53 signature, proliferative dormant and active STICs found amongst the cohort.

**Fig. S3.**
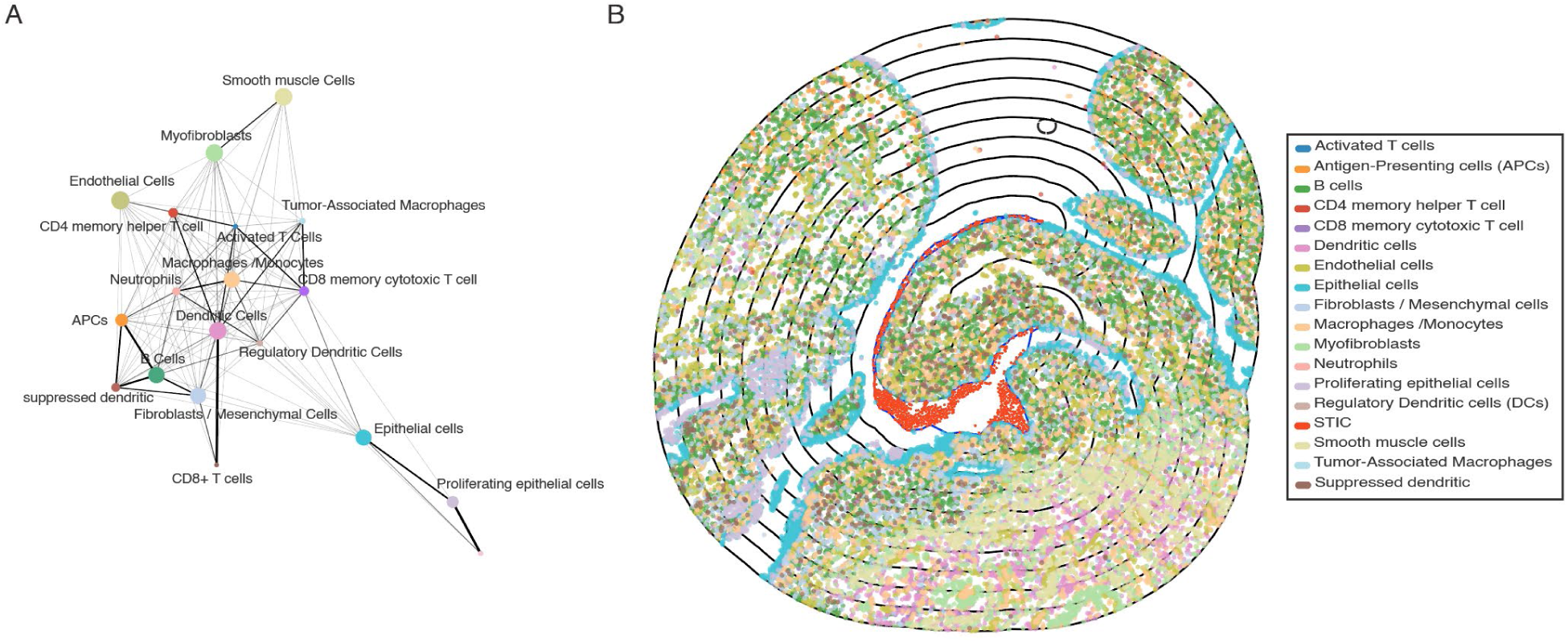
Topological and spatial proteomic mapping of proliferative active STIC microenvironment. **(a)** PAGA unpruned network analysis shows cellular interactions, highlighting immunosuppressive populations near proliferative active STIC. **(b)** Active STIC region was masked and subsequently dilated to generate 10 micron distances. At each distance, the cellular composition was assessed. Black lines in the figure indicate incremental 100 micron distance from the STIC in red.

**Fig. S4.**
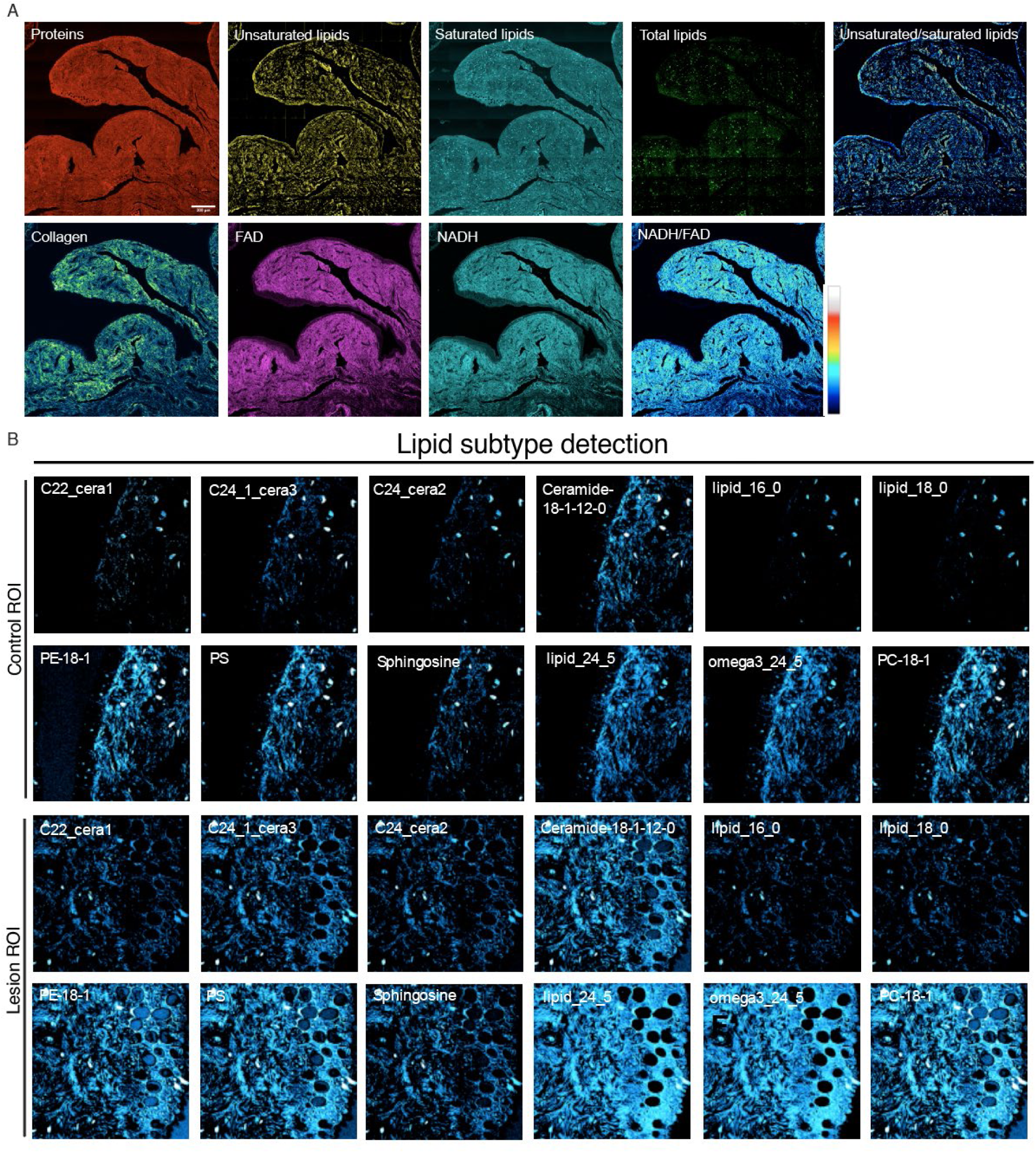
STIC lipid subtype characterization. **(a)** Multimodal SRS imaging displays the protein, lipid, unsaturated/saturated lipid, NADH, FAD and radiometric images of unsaturated/saturated lipids, NADH/FAD of the ROI shown in (Figure 1A). Scale bar: 200 µm. **(b)** SRS hyperspectra based lipid subtype detection showing the difference in lipid subtype levels between control and lesion ROI.

**Fig. S5.**
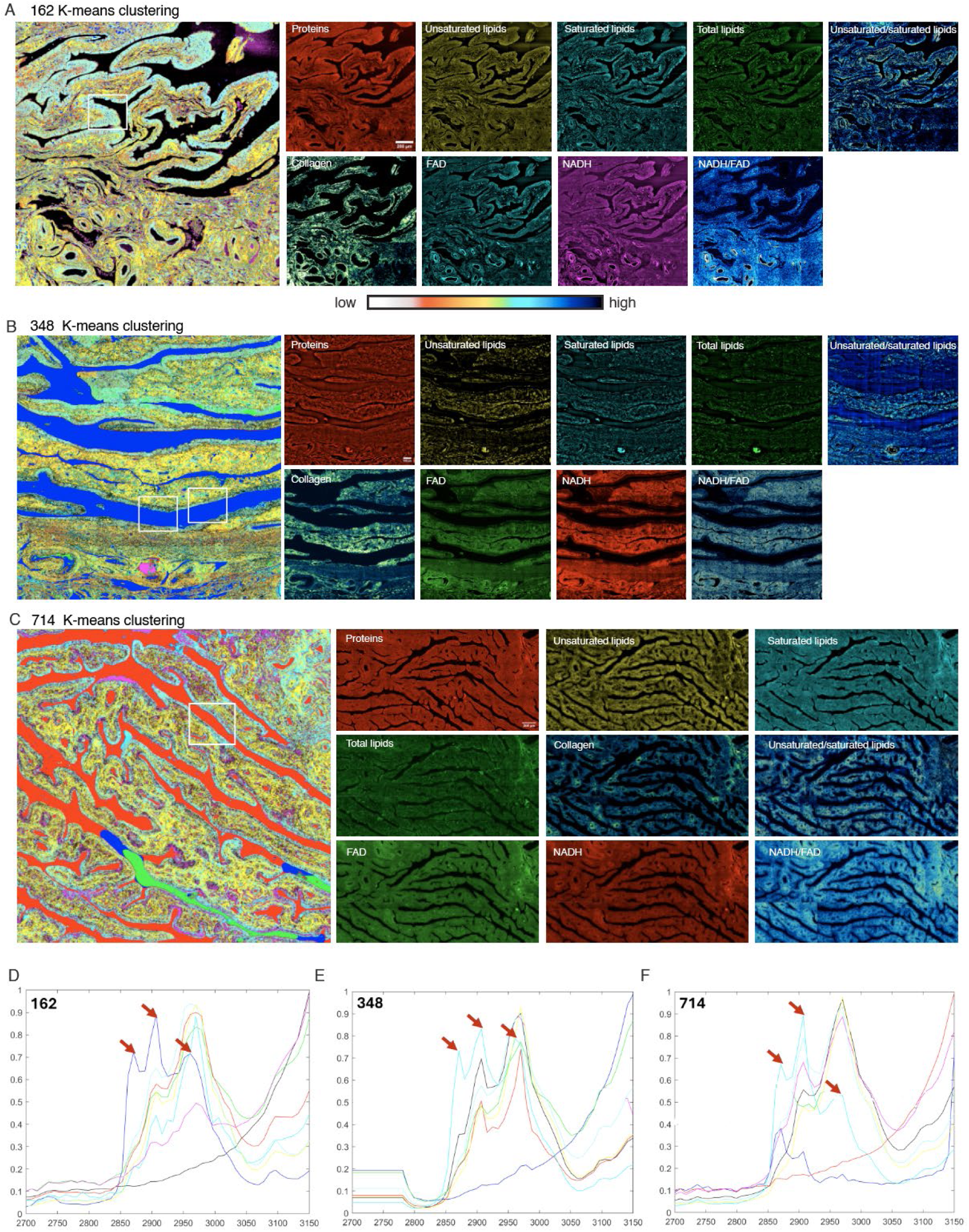
Spatial metabolomic profiling on additional targeted ovarian cancer lesions. **(a-c)** Multimodal and hyperspectral SRS imaging displays the metabolic states changes in multiple lesion tissues. Scale bar: 200 µm. **(d-e)** SRS hyperspectra from different lesion tissues underscore the consistent lipid metabolic remodeling. Red arrows points to the typic Raman lipid peaks at 2850 cm-1 and 2880 cm-1.

**Fig. S6.**
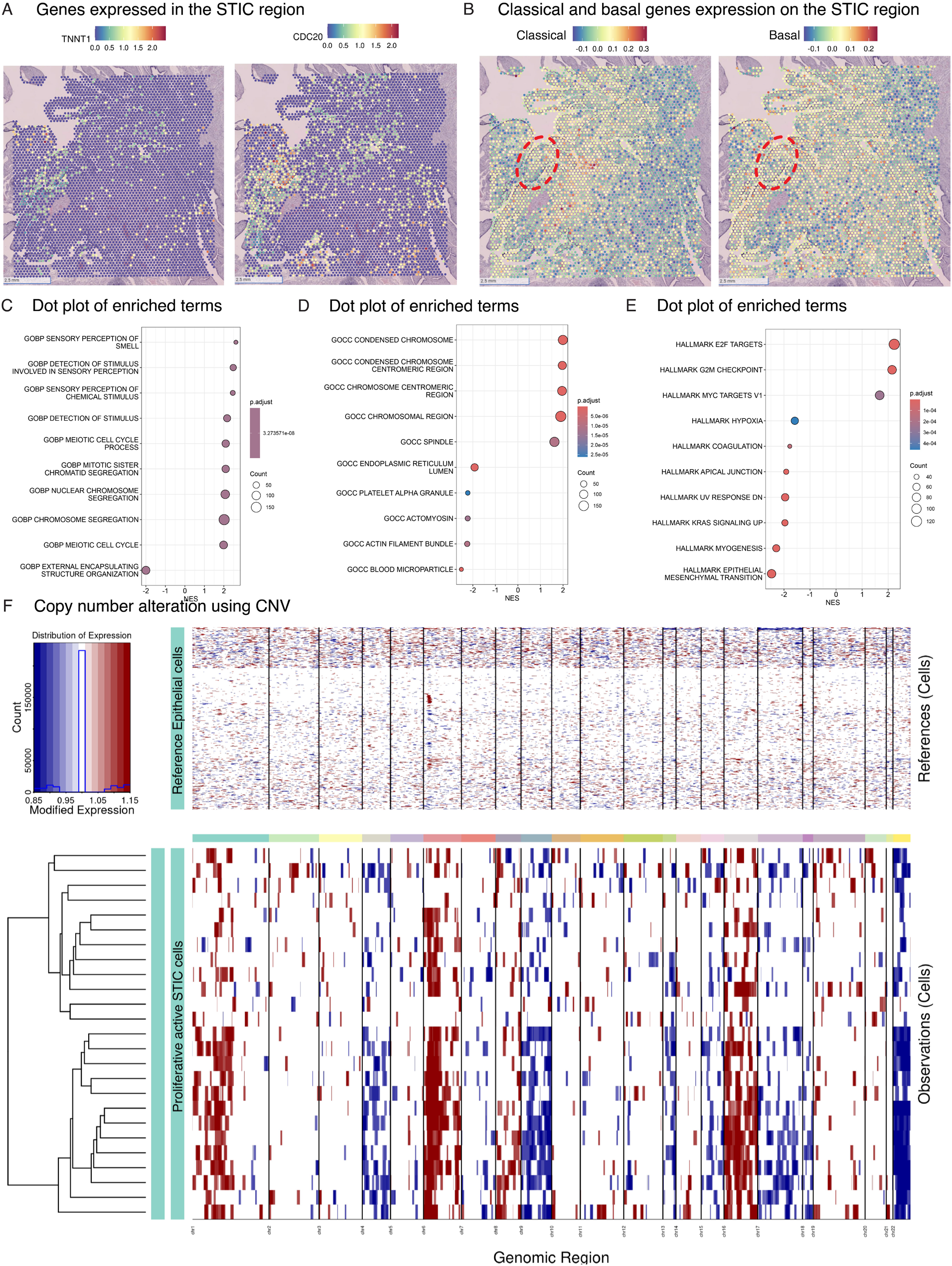
Spatial expression of ovarian cancer related genes, and pathway analysis. **(a)** TMNT1 and CDC20 gene expression patterns specific to STIC regions. **(b)** Sets of classical and basal gene expression did not show any STIC specific expression. **(c)** Dot plots showing significantly enriched terms from Gene Ontology Biological Process. **(d)** Dot plots showing significantly enriched terms from Gene Ontology Cellular Components. **(e)** Dot plots showing significantly enriched terms from Hallmark gene set. Dot size represents gene count, color intensity indicates adjusted p-value, and x-axis shows normalized enrichment score (NES). Terms are ordered by statistical significance. **(f)** Inferred CNA using inferCNV for two Visium sections of the same proliferative active STIC. Chromosomal gains are shown in red, and chromosomal depletions are shown in blue, with respect to reference adjacent healthy epithelial cells (top rows).

## SUPPLEMENTAL TABLES

**Table S1.**
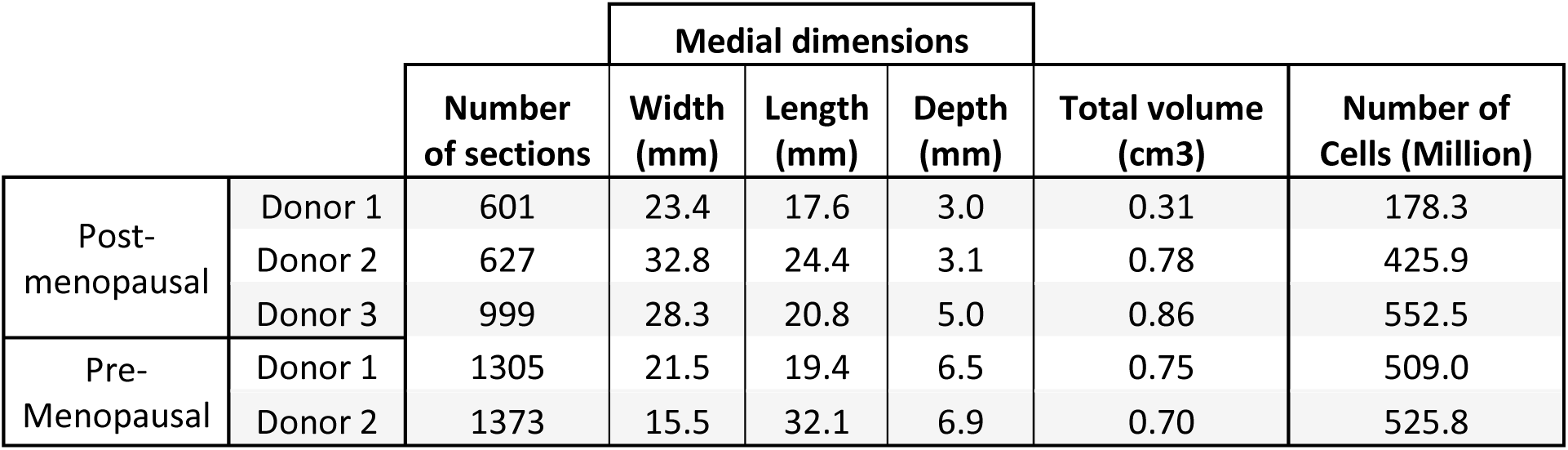
Table containing the number of Whole Slide Images, dimensions, and total volume of the whole human fallopian tube cohorts.

**Table S2.** Table containing the number of STICs found on the whole human fallopian tube cohorts.

|  |  | Precancerous lesions<br>(Proliferative active and dormant STICs) |  |  |  |  |
| --- | --- | --- | --- | --- | --- | --- |
|  |  | total | mean | median | min | max |
| Pre-menopausal | Donor 1 | 0 | 8.5 | 8.5 | 0 | 17 |
|  | Donor 2 | 17 |  |  |  |  |
| Post-menopausal | Donor 3 | 35 | 27.33 | 35 | 5 | 42 |
|  | Donor 4 | 42 |  |  |  |  |
|  | Donor 5 | 5 |  |  |  |  |

**Table S3.**
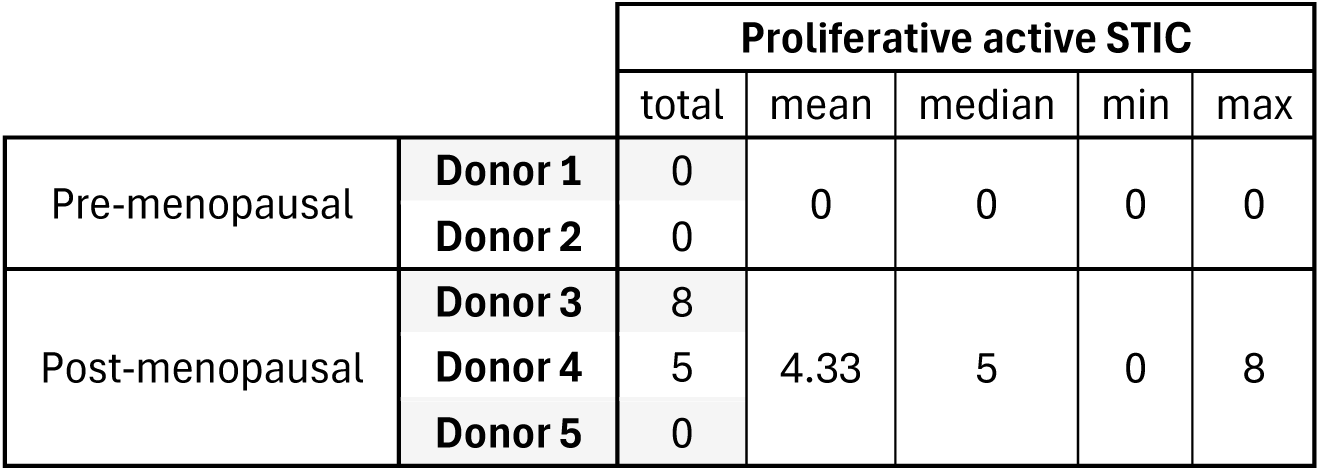
Table containing the number of STICs found on the whole human fallopian tube cohorts.

|  |  | Proliferative active STIC |  |  |  |  |
| --- | --- | --- | --- | --- | --- | --- |
|  |  | total | mean | median | min | max |
| Pre-menopausal | Donor 1 | 0 | 0 | 0 | 0 | 0 |
|  | Donor 2 | 0 |  |  |  |  |
| Post-menopausal | Donor 3 | 8 | 4.33 | 5 | 0 | 8 |
|  | Donor 4 | 5 |  |  |  |  |
|  | Donor 5 | 0 |  |  |  |  |

**Table S4.** Table containing the number of proliferative dormant STICs found on the whole human fallopian tube cohorts.

|  |  | Proliferative dormant STIC |  |  |  |  |
| --- | --- | --- | --- | --- | --- | --- |
|  |  | total | mean | median | min | max |
| Pre-menopausal | **Donor 1** | 0 | 8.5 | 8.5 | 0 | 17 |
|  | **Donor 2** | 17 |  |  |  |  |
| Post-menopausal | **Donor 3** | 27 | 23.00 | 27 | 5 | 37 |
|  | **Donor 4** | 37 |  |  |  |  |
|  | **Donor 5** | 5 |  |  |  |  |

**Table S5.** Table containing the number of p53 signatures found on the whole human fallopian tube cohorts.

|  |  | p53 signatures |  |  |  |  |
| --- | --- | --- | --- | --- | --- | --- |
|  |  | total | mean | median | min | max |
| Pre-menopausal | **Donor 1** | 1 | 3 | 3 | 1 | 5 |
|  | **Donor 2** | 5 |  |  |  |  |
| Post-menopausal | **Donor 3** | 1 | 1.67 | 1 | 0 | 4 |
|  | **Donor 4** | 4 |  |  |  |  |
|  | **Donor 5** | 0 |  |  |  |  |

**Table S6.** Table containing statistical prevalence of ovarian cancer precursor lesions in whole human fallopian tube cohorts.

| Group | # Patients | Proliferative active STIC Prevalence | Proliferative dormant STIC Prevalence | p53 Signature Prevalence |
| --- | --- | --- | --- | --- |
| Pre-menopausal | 2 | 0% (0/2) | 50% (1/2) | 50% (1/2) |
| Post-menopausal | 3 | 66.67% (2/3) | 100% (3/3) | 66.67% (2/3) |
| Combined | 5 | 40% (2/5) | 80% (4/5) | 60% (3/5) |

## METHODS

### Tissue acquisition and processing of entire human fallopian tubes

After resection of non-diseased human fallopian tubes from a donor network (nPOD). Specimens were processed into FFPE tissue blocks. Then, exhaustively serial sectioned at 4 microns and H&E stained (one every two sections). Unstained slides were stored in -20°C, under optimal humidity and vacuum conditions.

H&E-stained slides were scanned at 20x resolution (∼0.5 micron/pixel) using a Hamamatsu Nanozoomer S210. NDPI files were converted to tiff images (1 micron/pixel) and aligned into a 3D volume. StarDist method was employed to perform nuclear segmentation of all H&E-stained whole slide in entire fallopian tube samples.

### CODA microanatomical labelling of WSI of human fallopian tubes

To label the microanatomical components of the human fallopian tube, we developed two CODA semantic segmentation models^55,122^. One model labelled the surrounding epithelium microenvironment, including blood vessels, nerves, vasculature, mesothelium, rete ovarii in all WSI (Fig. S1, middle top panel). The second model was designed to automatically annotate the secretory and ciliated epithelial cells (Fig. S1, middle bottom panel). Models were combined to fully segmented all whole slide images in the human fallopian tubes (Fig. S1, right panel). InterpolAI was used to generate missing images to restore microanatomical connectivity^123^.

### Alignment of 2D WSI into 3D maps of entire human fallopian tubes

Combination of global rigid and local elastic image registrations allowed reconstruction of microanatomical structures of human fallopian tubes into 3D volume^55,61^. Alignment was applied to images subtyping the epithelium and to images labelling the fallopian tube microenvironment.

### Nuclear segmentation in H&E-stained images

To extract all 2.19 billion nuclear segmentations from 2,452 H&E-stained images, we used an adapted version of the StarDist pipeline for 3D histological slides (Fig. S1C)^124,125^. StarDist 40x resolution H&E segmentation pretrained model was finetuned to 20x resolution NDPI file format images^67^. To finetune the model, we annotated 25 H&E stained tiles with 256x256 dimensions for training. Training was optimized finetuning hyperparameters such as learning rate, training epochs, data augmentation. To maximize the heterogeneity of the testing tiles, we se4lected tiles from regions of the human fallopian tubes and across different specimens.

### Registration of 2D nuclear segmentation into 3D cellular volume

Similarly to the semantic segmentation step, segmented cell nuclei was registered into a 3D aligned volume, using CODA point cloud base registration method, which allowed the same alignment of the cell nuclei centroids accordingly with the tissue labelling registration^124^. Each cell on the 3D volume contained a unique cell ID, which allowed to link each cell to its respective morphological features.

### Measurement of bulk cellular and volumetric quantifications

With the generated 3D tissue and cellular volumes, bulk quantifications can be extracted. To obtain volumetric data from each respective label, all voxels of each respective tissue component are summed and, subsequently, adjusted according to its respective voxel size. Bulk cellular information of each microanatomical label can be extracted *in silico* by combination of 3D tissue labelled volume with its respective label locations in the 3D cellular volume.

### 3D cellular and volumetric variability within human fallopian tube epithelium

To quantify the variability in cellular and volumetric content within each human fallopian tube, a virtual path was generated along the epithelium. Along this virtual epithelium path, cross sections perpendicular to this path were generated to simulate travelling across the human fallopian tube epithelium. Cellular and volumetric measurements were obtained for each cross section, resulting in tens of thousands of virtual cross sections along each fallopian tube.

### Detection of p53 signatures, proliferative dormant and active STICs in human fallopian tubes

To 3D map precursors to ovarian cancer in whole human fallopian tubes, we developed a framework that integrates H&E and IHC (*p53* and *Ki67*) staining methods in 3D. First, we developed a deep learning method to identify positive signal locations in *p53* and *Ki67* IHC stained images (stained one in every 8 sections of the entire stack of images). Then we aligned the IHC images to the aligned H&E-stained image stack. Combination of IHC stained slides and H&E-stained slides to highlight regions with potential precursors of ovarian cancer. Generation of the lesions image stacks containing IHC and H&E images to manually check hundreds of potential lesion locations across different specimens. Manual validation of each bounding box with pathologists and trained experts allowed the compiling of confirmed ovarian cancer precursors. Validated lesions were then used for further multi-omics profiling.

### Volume distribution of ovarian cancer precursors across different samples

For each individual validated lesion, their volume was computed (Fig. 3F, left panel). Power law was used to predict lesion growth for proliferative active and dormant STICs, and p53 signatures combined (Fig. 3F, middle panel)^65^. Using the Kolmogorov-Smirnov test, maximization of the p-value was applied to fit the measured lesion volumes^126^.

### 3D virtual SEE-FIM procedure for detection of epithelial lesions in low-risk nondiseased human fallopian tube samples

To virtually simulate the SEE-FIM procedure in our samples, we generated a virtual path across the human fallopian tube’s epithelium. In the fimbriated ends of the fallopian tube, longitudinal sections were generated, and on the remaining of the fallopian tube transverse cross sections were computed. For each fallopian tube sample, equally distanced slides were generated ranging from 1 up to 11,000 sections along the epithelium’s center path (Fig. 3G, and Fig. S2D). Simulations of the distinct virtual section ranges were computed for each fallopian tube (Fig. S2E) and, for each combination of sections simulated, the number of lesions was assessed. The same was computed to the respective percentage of the fallopian tube sectioned (Fig. 3H).

### Spatial proteomics on region of interest to map STIC immune landscape

To deeply profile the proteomic landscape involved in STIC progression, we applied CODEX spatial proteomics using 25 marker antibody panel targeting epithelial, immune, and stromal populations (Fig. 4A). WSI cyclic immunofluorescence was conducted to ensure spatial comparison of STIC to non-lesional epithelia. DAPI nuclear channel was segmented and subsequent dilation of the nuclear area ensured cells were isolated and boundaries of each were accurately delimited (Fig. 4B)^67^. For each segmented cell, protein expression intensities were quantified. Marker intensities were normalized to minimize the effects of inter-cell variability. UMAP was applied to visualize multidimensional protein expression profiles and identify distinct cellular clusters (Fig. 4C)^68–71,127–133^. Clusters were annotated based on canonical marker expressions to distinguish epithelial, stromal, and immune cell populations (Fig. 4D-F).

Spatial mapping was then performed to measure the distribution of cell types across the STIC and adjacent epithelial regions (Fig. 4G-I). To investigate the cell-to-cell interaction between the cell phenotypes, we applied a PAGA (Partition-based Graph Abstraction)^72^. Using PAGA, we inferred proximity and connectivity between cell phenotypes (Fig. 4G, S3A).

To compute the variance in cellular composition relative to STIC proximity, we performed a spatial dilation of the STIC mask and measured the cellular composition at different distances (Fig. S3B). Cell type composition was calculated every 10 µm distances and extended up to 500 µm from STIC region (Fig. 4H). Quantification of the differential cellular enrichment was quantified by comparing regions within 100 µm to this STIC to regions distancing 400 and 500 µm from the STIC (Fig. 4I).

### Two-photon stimulated Raman scattering to map metabolomic signature in STIC

Pathologist-validated lesions within 3D spatial maps of fallopian tubes were selected for spatial metabolomics analysis of the ovarian cancer precursors (Fig. 5A). High-resolution multimodal two-photon stimulated Raman scattering (SRS) imaging was used to quantify metabolic signatures across lesions^59^. Multimodal SRS imaging targeting lesions and control normal adjacent epithelial regions of interest (ROIs) allowed to capture spatial distributions of proteins, saturated and unsaturated lipids, total lipids, NADH, and FAD (Fig. 5B-C). Intensity-normalized images were concatenated into feature vectors and analyzed by principal-component analysis followed by k-means clustering (Fig. 5D-E)^134^.

**Fig. 5.**
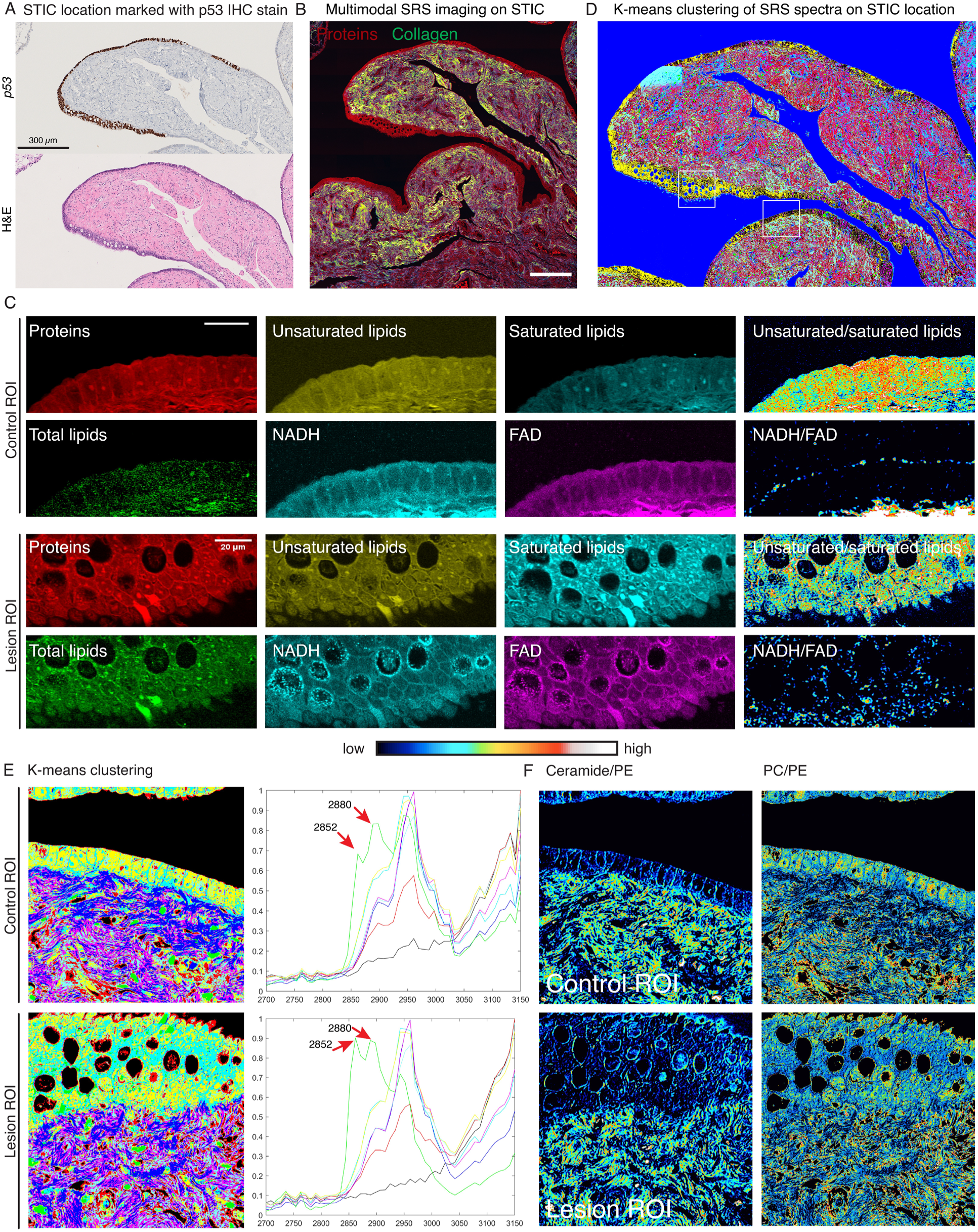
Multimodal SRS imaging reveals the metabolic remodeling in lesions. **(a)** P53 staining shows the lesion regions. **(b)** SRS protein channel overlayed with second homogenization (SHG) signal for collagen to show the architecture of the same lesion region of interest (ROI) in (A). Scale bar: 200 µm. **(c)** Multimodal SRS imaging shows the comparation of multiple metabolites between control and lesion ROI. Scale bar: 20 µm. **(d)** Hyperspectral SRS imaging and unsupervised clustering showing the distribution of metabolites across the whole ROI shown in (A) and (B). **(e)** Hyperspectral SRS imaging and unsupervised clustering shows the difference of metabolic profiles between control and lesion ROI. Red arrows points to the typic Raman lipid peaks at 2850 cm^-1^ and 2880 cm^-1^. **(f)** The ratio images of ceramide/PE and PC/PE highlight the lipid subtype modulation between control and lesion ROI.

### Spatial transcriptomic profiling of low-risk non-diseased STIC

3D mapped regions containing STIC were selected and used to guide placement of the Visium CytAssist capture area (6.5 x 6.5 mm^2^). Tissue sections were processed using the 10x Genomics Visium CytAssist FFPE protocol. After deparaffinization and epitope retrieval, hybridization with the Human Transcriptome Probe Set v2.0. Probe release was conducted via CytAssist, followed by library preparation and sequencing (∼250 million reads/sample) on an Illumina NovaSeq 6000^121,135^.

Differential gene expression analysis between STIC and normal regions was performed using Seurat v5 ^136^. Genes with adjusted p-value <0.05 and log_2_ fold change >0.25 were considered significant (Fig. 6C, S6A-B). Significantly altered genes were subjected to Gene Ontology (GO) analysis (Fig. 6E) and Hallmark pathway enrichment using GSEA (Fig. 6F–G, S6C-E)^137–139^. Enrichment scores were visualized using dot plots and ranked enrichment plots. Copy number alterations were inferred from transcriptomic data using inferCNA and inferCNF, comparing STIC spots to adjacent normal epithelial reference regions^60,140^. Alterations were visualized as heatmaps and chromosome-specific dot plots (Fig. 6H–I). Spatial distribution of chromosomal alterations was mapped across tissue sections (Fig. 6J–K, Fig. S6F).

**Fig. 6.**
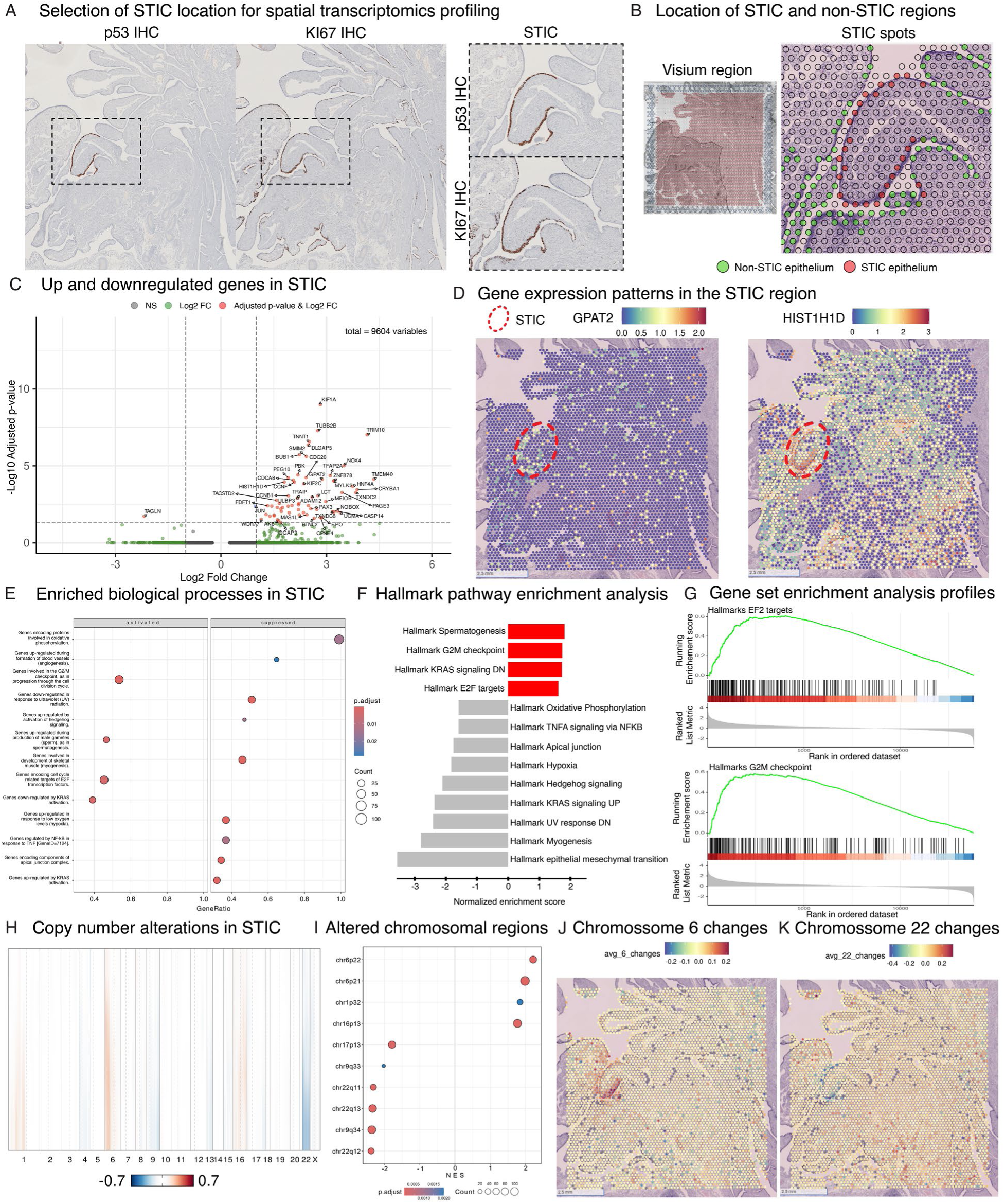
Spatial transcriptomic profiling reveals molecular alterations in STIC lesions. **(a)** Selection of STIC regions for spatial transcriptomics profiling, validated by *p53* and *Ki67* IHC. **(b)** Identification of STIC and non-STIC epithelial regions within the Visium spatial transcriptomics platform. **(c)** Heatmap of differentially expressed genes in STIC lesions, highlighting key upregulated and downregulated targets (adjusted p-value < 0.05). **(d)** Oncogene expression patterns (e.g., *GPAT2*, *HIST1H1D*) specific to STIC regions. **(e-f)** Dot plots of enriched gene signatures in STIC, including KRAS signaling, oxidative phosphorylation, and epithelial-mesenchymal transition. **(g)** Chromosomal ploidy analysis showing copy number variations in STIC, with focal changes on chromosomes 6 and 22.

### Statistical considerations

All significance tests were performed using the Wilcoxon rank sum test. To compare metrics within and between cohorts, median, mean, standard deviation, and interquartile range were determined. Relative error was defined as [measured value – expected value] / expected value. No other statistical calculations were performed in this work.

## Code and data availability statement

The data analyzed here is available from the corresponding author upon request. The 3D volumes created here will be made publicly available on Zenodo upon publication of this manuscript. The code used to generate the 3D tissue maps is available on the following GitHub: https://github.com/ashleylk/CODA. The code to develop the 3D maps of precancerous lesions will be made available upon publication.

## Author contributions

A.F., A.L.K. and D.W. conceived the project. A.F., A.C., V.Q., B.P., R.O.F., M.A., P. G., M. B. collected and processed the human fallopian tube samples. A.F., A.L.K, V.Q, Y.L, A.H, M.T., S.J, D.K, O.N., H.B., D.Z., L.K., L.S., R.F. and P.W. conducted the analysis. I.M.S validated histological analysis as clinical experts in pancreatic cancer pathology. A.F., A.L.K., and D.W. wrote the first draft of the manuscript, which all authors edited and approved.

